# A lytic transglycosylase connects bacterial focal adhesion complexes to the peptidoglycan cell wall

**DOI:** 10.1101/2024.04.04.588103

**Authors:** Carlos A. Ramírez Carbó, Olalekan G. Faromiki, Beiyan Nan

## Abstract

The Gram-negative bacterium *Myxococcus xanthus* glides on solid surfaces. Dynamic bacterial focal adhesion complexes (bFACs) convert proton motive force from the inner membrane into mechanical propulsion on the cell surface. It is unclear how the mechanical force transmits across the rigid peptidoglycan (PG) cell wall. Here we show that AgmT, a highly abundant lytic PG transglycosylase homologous to *Escherichia coli* MltG, couples bFACs to PG. Coprecipitation assay and single-particle microscopy reveal that the gliding motors fail to connect to PG and thus are unable to assemble into bFACs in the absence of an active AgmT. Heterologous expression of *E. coli* MltG restores the connection between PG and bFACs and thus rescues gliding motility in the *M. xanthus* cells that lack AgmT. Our results indicate that bFACs anchor to AgmT-modified PG to transmit mechanical force across the PG cell wall.

## Introduction

In natural ecosystems, the majority of bacteria attach to surfaces^1^. Surface-associated motility is critical for many bacteria to navigate and populate their environments^2^. The Gram-negative bacterium *Myxococcus xanthus* moves on solid surfaces using two independent mechanisms: social (S-) motility and adventurous (A-) motility. S-motility, analogous to the twitching motility in *Pseudomonas* and *Neisseria*, is powered by the extension and retraction of type IV pili^3,4^. A-motility is a form of gliding motility that does not depend on conventional motility-related cell surface appendages, such as flagella or pili^2,5^. AglR, AglQ, and AglS form a membrane channel that functions as the gliding motor by harvesting proton motive force^6,7^. Motor units associate with at least fourteen gliding-related proteins that reside in the cytoplasm, inner membrane, periplasm, and outer membrane^8,9^.

In each gliding cell, two distinct populations of gliding complexes coexist: dynamic complexes move along helical paths and static complexes remain fixed relative to the substrate^10–13^. Motors transport incomplete gliding complexes along helical tracks but form complete, force-generating complexes at the ventral side of cells that appear fixed to the substrate (**Fig. 1**)^10,13^. Based on their functional analogy with eukaryotic focal adhesion sites, these complete gliding machineries are called bacterial focal adhesion complexes (bFACs). Despite their static appearance, bFACs are dynamic under single-molecule microscopy. Individual motors frequently display move-stall-move patterns, in which they move rapidly along helical trajectories, join a bFAC where they remain stationary briefly, then move rapidly again toward the next bFAC or reverse their moving directions^11,14,15^. bFACs adhere to the gliding substrate through an outer membrane adhesin^12^. As motors transport bFACs toward lagging cell poles, cells move forward but bFACs remain static relative to the gliding substrate.

**Fig. 1.**
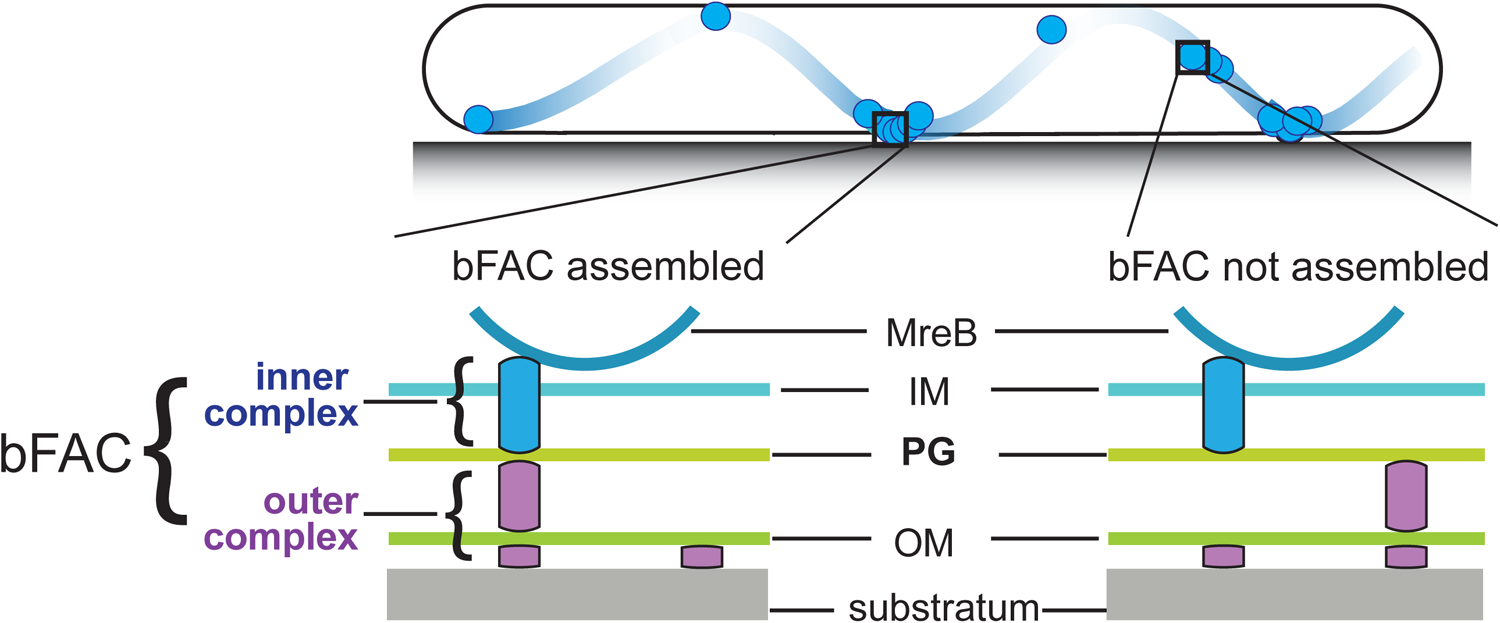
Stationary bFACs drive *M. xanthus* gliding. Motors carrying incomplete gliding complexes either diffuse or move rapidly along helical paths but do not generate propulsion. Motors stall and become nearly static relative to the substrate when they assemble into complete bFACs with other motor-associated proteins at the ventral side of the cell. Stalled motors push MreB and bFACs in opposite directions and thus exert force against outer membrane adhesins. Overall, as motors transport bFACs toward lagging cell poles, cells move forward but bFACs remain static relative to the substrate. IM, inner membrane; OM, outer membrane.

bFAC assembly can be quantified at nanometer resolution using the single-particle dynamics of gliding motors^11,16^. Single particles of a fully functional, photoactivatable mCherry (PAmCherry)-labeled AglR display three dynamic patterns, stationary, directed motion, and diffusion^11,16^. The stationary population consists of the motors in fully assembled bFACs, which do not move before photobleach. In contrast, the motors moving in a directed manner carry incomplete gliding complexes along helical tracks, whereas the diffusing motors assemble to even less completeness^10,11,13,16^. As motors only generate force in static bFACs^10^, the population of nonmotile motors indicates the overall status of fully assembled bFACs.

However, how bFACs transmit force across the peptidoglycan (PG) cell wall is unclear. PG is a mesh-like single molecule of crosslinked glycan strands that surrounds the entire cytoplasmic membrane. The rigidity of PG defines cell shape and protects cells from osmotic lysis^17,18^. If bFACs physically penetrate PG, their transportation toward lagging cell poles would tear PG and trigger cell lysis. To avoid breaching PG, an updated model proposes that rather than forming stable and rigid complexes, gliding-related proteins only assemble into force-generating machineries in bFACs.

Outside of bFACs, these proteins could localize diffusively or move along with unengaged motor units^10,19^. To simplify this model, we can artificially divide each gliding machinery into two subcomplexes, one on each side of the PG layer. The inner subcomplexes move freely and only assembly with the outer subcomplexes in bFACs, which transform proton motive force into mechanical propulsion on cell surfaces (**Fig. 1**).

The inner subcomplexes, containing the motors, are the force-generating units in bFACs. As motors reside in the fluid inner membrane, to transmit force to the cell surface, the inner subcomplexes must push against two relatively rigid structures, one on each side of the membrane, in opposite directions^19^ (**Fig. 1**). MreB is a bacterial actin homolog that supports rod shape in many bacteria^16,17,20^. In *M. xanthus*, MreB also connects to bFACs on the cytoplasmic side and thus plays essential roles in gliding^2,5,6,12,16,21,22^. The inner subcomplexes could push against MreB filaments and PG in the cytoplasm and periplasm, respectively^10,16,19,23^. The interaction between the inner subcomplexes and PG not only satisfies the physical requirement for force generation but also supports bFAC stability. Without this interaction, the inner and outer subcomplexes can only form transient, “slippery” association, which is predicted to produce short and aberrant cell movements^19^. How the inner complex interacts with PG remains unknown. It was speculated that gliding motors in the inner complex could bind PG directly^10^. However, such binding has not been confirmed by experiments.

In this study, we found that AgmT, a lytic transglycosylase (LTG) for PG, is required for *M. xanthus* gliding motility. Whereas AgmT only regulates cell morphology moderately during vegetative growth, it is essential for maintaining PG integrity under the stress from the antibiotic mecillinam. Using single-particle tracking microscopy and coprecipitation assays, we found that AgmT is essential for the inner subcomplexes to connect to PG and stall in bFACs. Importantly, expressing *Escherichia coli* MltG heterologously rescues the connection between PG and bFACs and thus restores gliding motility. Hence, the LTG activity of AgmT anchors bFACs to PG, potentially by modifying PG structure. Our findings reveal the long-sought connection between PG and bFACs that allows mechanical force to transmit across the PG cell wall.

## Results

### AgmT, a putative LTG, is required for gliding motility

To elucidate how bFACs interact with PG, we searched for potential PG-binding domains among the proteins that are required for gliding motility. A previous report identified 35 gliding-related genes in *M. xanthus* through transposon-mediated random mutagenesis^24^. Among these genes, *agmT* (ORF K1515_04910^25^) was predicted to encode an inner membrane protein with a single transmembrane helix (residues 4 – 25) followed by a large “periplasmic solute-binding” domain^24^. No other motility-related genes are found in the vicinity of *agmT*. After careful analysis, we found that AgmT showed significant similarity to the widely conserved YceG/MltG family LTGs. The putative active site, Glu223 (corresponding to E218 in *E. coli* MltG)^26^, is conserved in AgmT (**Fig. 2A**).

**Fig. 2.**
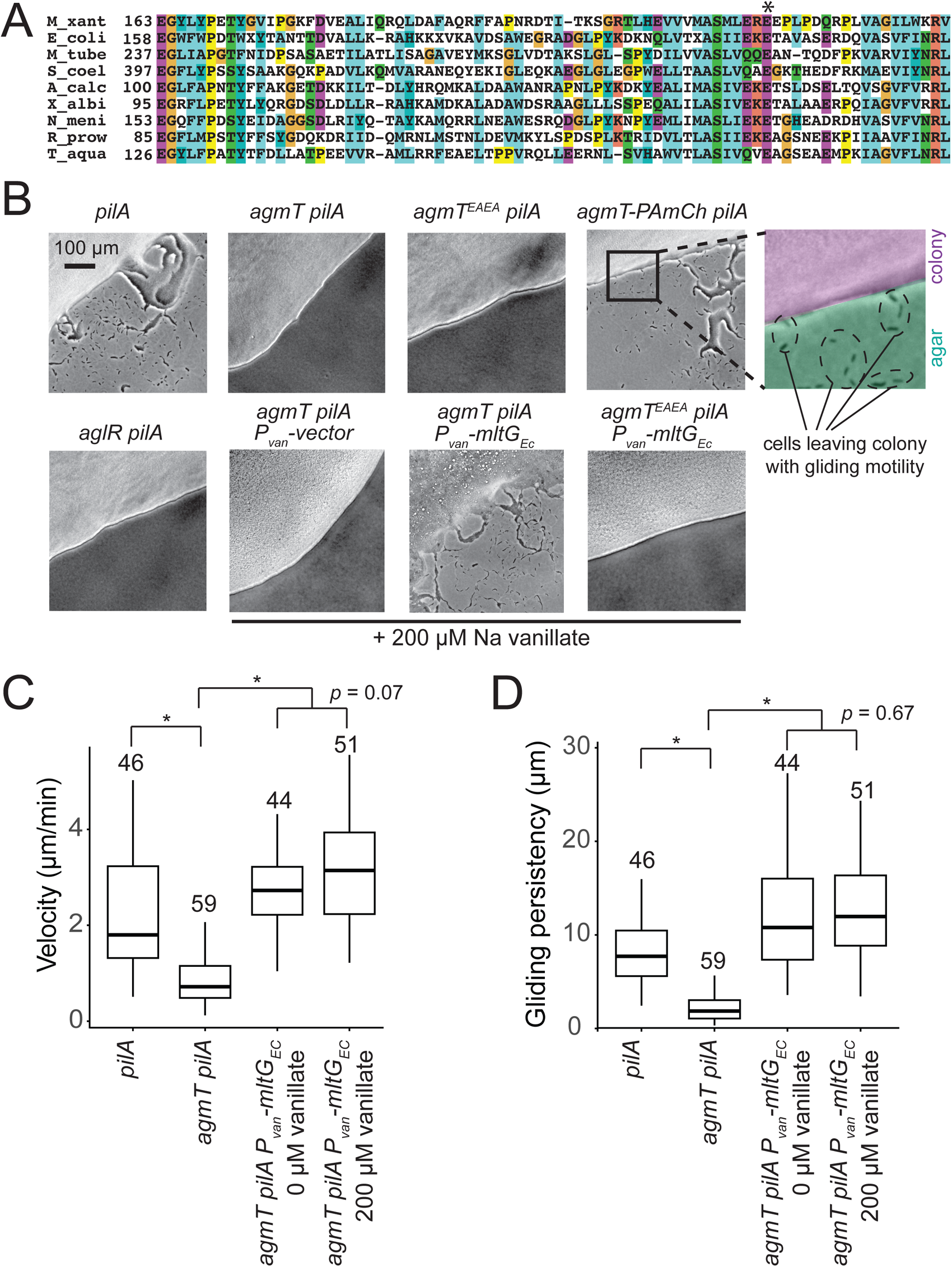
AgmT, a putative lytic transglycosylase, is required *for M. xanthus* gliding. **A)** AgmT shows significant similarity to a widely conserved PG transglycosylase in YceG/MltG family. The conserved glutamine residue is marked by an asterisk. M_xant, *M. xanthus*; E_coli, *E. coli*; M_tube, *Mycobacterium tuberculosis*; S_coel, *Streptomyces coelicolor*; A_calc, *Acinetobacter calcoaceticus*; X_albi, *Xanthomonas albilineans*; N_meni, *Neisseria menningitidis*; R_prow, *Rickettsia prowazekii*; T_aqua, *Thermus aquaticus*. **B)** AgmT is required for *M. xanthus* gliding. Colony edges were imaged after incubating cells on 1.5% agar surface for 24 h. To eliminate S-motility, we further knocked out the *pilA* gene that encodes pilin for type IV pilus. Cells that move by gliding are able to move away from colony edges. Deleting *agmT* or disabling the active site of AgmT abolish gliding but fusing an PAmCherry (PAmCh) to its C-terminus does not. Heterologous Expression of *E. coli* MltG (MltGEc) restores gliding of *agmT* cells but not the cells that express AgmT^EAEA^. **C, D)** While cells lacking AgmT moved slower **(C)** and less persistently (**D**, measured by the distances cells traveled before pauses and reversals), the expression of MltGEc restores both the velocity and persistence of gliding in the *agmT* cells. Data were pooled from three independent experiments and *p* values were calculated using a one-way ANOVA test between two unweighted, independent samples. Boxes indicate the 25^th^ - 75^th^ percentiles and bars the median. The total number of cells analyzed is shown on top of each plot. *, *p* <0.001.

To confirm the function of AgmT in gliding, we constructed an *agmT* in-frame deletion mutant. We further knocked out the *pilA* gene that encodes pilin for type IV pilus to eliminate S-motility. On a 1.5% agar surface, the *pilA^-^* cells moved away from colony edges both as individuals and in “flare-like” cell groups, indicating that they were still motile with gliding motility. In contrast, the ***Δ****aglR pilA^-^* cells that lack an essential component in the gliding motor were unable to move outward from the colony edge and thus formed sharp colony edges. Similarly, the ***Δ****agmT pilA^-^* cells also formed sharp colony edges, indicating that they could not move efficiently with gliding (**Fig. 2B**).

We then imaged individual ***Δ****agmT pilA^-^* cells on 1.5% agar surface at 10-s intervals using bright-field microscopy. To our surprise, instead of being static, individual ***Δ****agmT pilA^-^* cells displayed slow movements, with frequent pauses and reversals (**Video 1**). To quantify the effects of AgmT, we measured the velocity and gliding persistency (the distances cells traveled before pauses and reversals) of individual cells. Compared to the *pilA^-^* cells that moved at 2.30 ± 1.33 µm/min (n = 46) and high persistency (**Video 2** and **Fig. 2C, D**), ***Δ****agmT pilA^-^* cells moved significantly slower (0.88 ± 0.62 µm/min, n = 59) and less persistent (**Video 1** and **Figure. 2C, D**). Such aberrant gliding motility is distinct from the “hyper reversal” phenotype due to the polarity regulators constitutively switching the moving direction of cells, because hyper reversing cells usually maintain gliding velocity at the wild-type level^27^. Instead, such phenotypes match the prediction that when the inner complexes of bFACs lose connection with PG, bFACs can only generate short, and inefficient movements^19^. Our data indicate that AgmT is not essential component in the bFACs. Thus, AgmT is likely to regulate the assembly and stability of bFACs, especially their connection with PG.

### AgmT is an LTG

As AgmT is a putative LTG for PG, we then tested if its predicted active site, Glu223, is required for gliding. Because the amino acid following Glu223 is also a glutamate (Glu224), we replaced both glutamate residues with alanine (AgmT^EAEA^) using site-directed mutagenesis to alter their codons on the chromosome. *agmT^EAEA^ pilA^-^* cells formed sharp colony edges on agar surface (**Fig. 2B**). Thus, the putative LTG activity of AgmT is required for *M. xanthus* gliding motility.

To determine if AgmT is an LTG, we expressed the periplasmic domain (amino acids 25 – 339) of wild-type AgmT and AgmT^EAEA^ in *E. coli*. We purified PG from wild-type *M. xanthus* cells, labeled it with Remazol brilliant blue (RBB), and tested if the purified AgmT variants hydrolyze labeled PG *in vitro* and release the dye^28,29^. Similar to lysozyme that specifically cleaves the β-1,4-glycosidic bonds in PG, wild-type AgmT solubilized dye-labeled *M. xanthus* PG that absorbed light at 595 nm. In contrast, the AgmT^EAEA^ variant failed to release the dye (**Fig. 3A**). Hence, AgmT displays LTG activity *in vitro*.

**Fig. 3.**
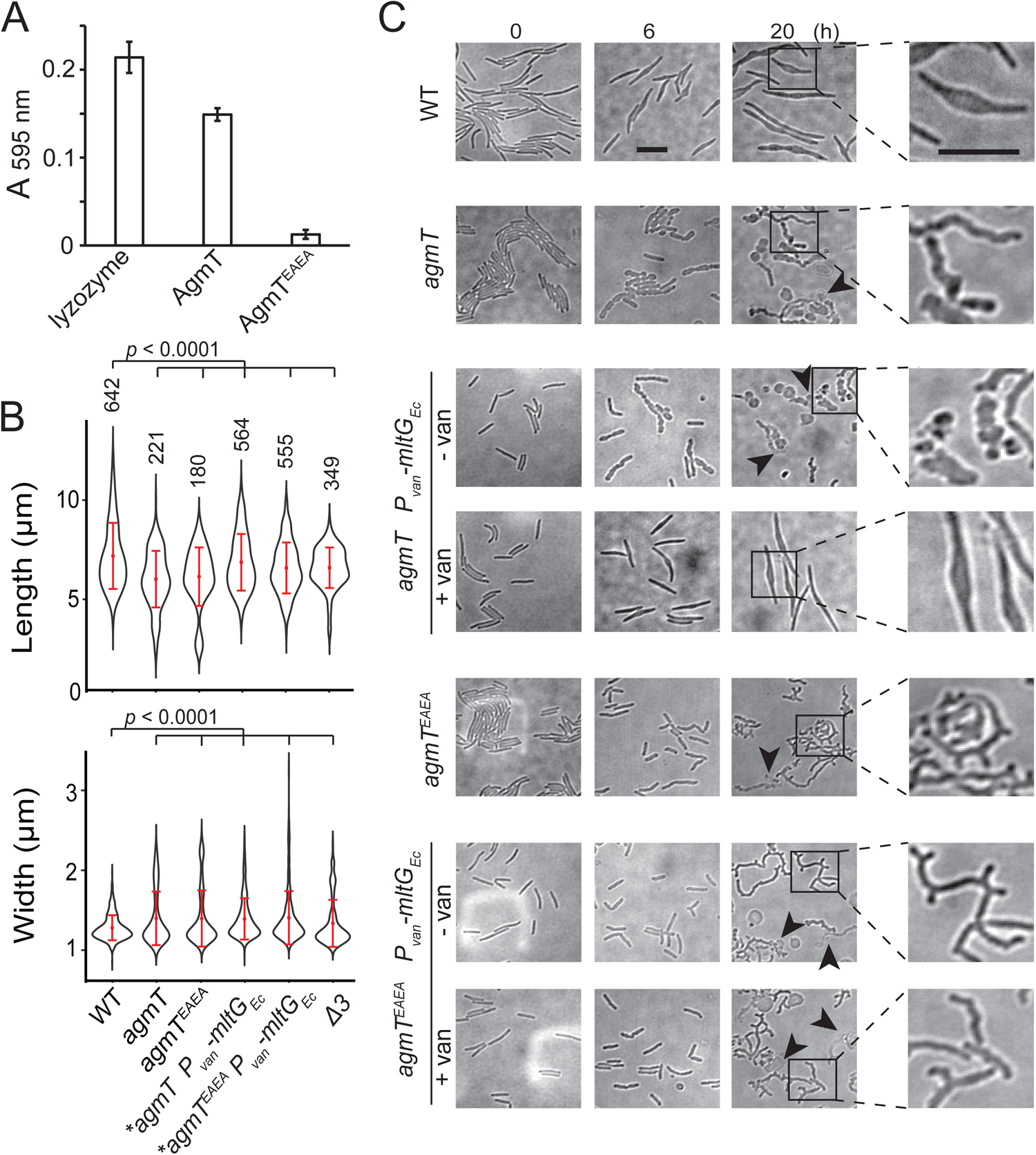
AgmT regulates cell morphology and integrity under antibiotic stress. A) Purified AgmT solubilizes dye-labeled PG sacculi, but AgmT^EAEA^ does not. Lysozyme serves as a positive control. Absorption at 595 nm was measured after 18 h incubation at 25 C. Data are presented as mean values ± SD from three replicates. **B)** AgmT regulates cell morphology. Compared to wild-type cells, cells that lack AgmT and express AgmT^EAEA^ are significantly shorter and wider. A previously reported mutant that lacks all three class A penicillin-binding proteins (*Δ3*) displays similarly shortened and widened morphology but is still motile by gliding (**Fig. S1**). Heterologous expression of *E. coli* MltG partially restores cell length but not cell width in *agmT* and *agmT^EAEA^* backgrounds. Asterisks, 200 M sodium vanillate added. *p* values were calculated using a one-way ANOVA test between two unweighted, independent samples. Whiskers indicate the 25^th^ - 75^th^ percentiles and red dots the median. The total number of cells analyzed is shown on top of each plot. **C)** AgmT regulates cell morphology and integrity under mecillinam stress (100 μg/ml). Expressing *E. coli* MltG by a vanillate-inducible promoter (*Pvan*) restores resistance against mecillinam in *agmT* cells but not in the cells that express AgmT^EAEA^. van, 200 M sodium vanillate. Arrows point to newly lysed cells. Scale bars, 5 µm.

Whereas AgmT does not affect growth rate (**Fig. S2**), cells that lacked AgmT or expressed AgmT^EAEA^ maintained rod shape but were slightly shorter and wider than the wild-type ones (**Fig. 3B**). Nevertheless, altered morphology alone does not likely account for abolished gliding. In fact, a previously reported mutant that lacks all three class A penicillin-binding proteins (*Δ3*) displays similarly shortened and widened morphology^30^ but is still motile by gliding (**Fig. S1**).

A recent report revealed that *Vibrio cholerae* MltG degrades un-crosslinked PG turnover products and prevents their detrimental accumulation in the periplasm^31^. To test if AgmT plays a similar role in *M. xanthus*, we used mecillinam to induce cell envelope stress. Mecillinam is a β-lactam that induces the production of toxic, un-crosslinked PG strands^32^. Rather than collapsing the rod shape, mecillinam only causes bulging near the centers of wild-type *M. xanthus* cells, whereas large-scale cell lysis does not occur, and cell poles still maintain rod shape even after prolonged (20 h) treatment (**Fig. 3C**)^30^. Thus, wild-type *M. xanthus* can largely mitigate mecillinam stress. In contrast, mecillinam-treated *agmT* cells increased their width drastically, displayed significant cell surface irregularity along their entire cell bodies, and lysed frequently (**Fig. 3C**). Cells expressing AgmT^EAEA^ as the sole source of AgmT displayed similar, but even stronger phenotypes under mecillinam stress, growing into elongated, twisted filaments that formed multiple cell poles and lysed frequently (**Fig. 3C**). These results confirm that similar to *V. cholerae* MltG, *M. xanthus* AgmT is important for maintaining cell integrity under mecillinam stress. The different responses from *agmT* and *agmT^EAEA^* cells suggest that other LTGs could partially substitute AgmT while AgmT^EAEA^ blocks these enzymes from accessing un-crosslinked PG strands.

AgmT is the only LTG *in M. xanthus* that belongs to the YceG/MltG family. Besides *agmT*, the genome of *M. xanthus* contains 13 genes that encode putative LTGs (**Table S1**). To test if these proteins also contribute to gliding, we knocked out each of these 13 genes in the *pilA^-^* background. The resulting mutants all retained gliding motility, indicating that AgmT is the only LTG that is required for *M. xanthus* gliding (**Fig. S1**).

### AgmT is essential for proper bFACs assembly

How does AgmT support gliding? We first tested if AgmT regulates the function of the gliding motor. For this purpose, we quantified the motor dynamics by tracking single particles of AglR, an essential component of the motor. We spotted the cells that express a fully functional, photo-activatable mCherry (PAmCherry)-labeled AglR^11^ on 1.5% agar surface. We used a 405-nm excitation laser to activate the fluorescence of a few labeled AglR particles randomly in each cell and quantified their localization using a 561-nm laser at 10 Hz using single particle tracking photo-activated localization microscopy (sptPALM) under highly inclined and laminated optical sheet (HILO) illumination^11,16,33^. Using this setting, only a thin section of each cell surface that was close to the coverslip was illuminated. To analyze the data, we only chose the fluorescent particles that remained in focus for 4 - 12 frames (0.4 - 1.2 s). As free PAmCherry particles diffuse extremely fast in the cytoplasm, entering and exiting the focal plane frequently, they usually appear as blurry objects that cannot be followed at 10-Hz close to the membrane^16^. For this reason, the noise from any potential degradation of AglR- PAmCherry was negligible. Consistent with our previous results^11,14^, 32.1% (n = 2,700) of AglR-PAmCherry particles remained within one pixel (160 nm × 160 nm) before photobleach, indicating that they were immotile. The remaining 67.9% AglR-PAmCherry particles were motile, leaving trajectories of various lengths (**Fig. 4A, 4B**).

**Fig. 4.**
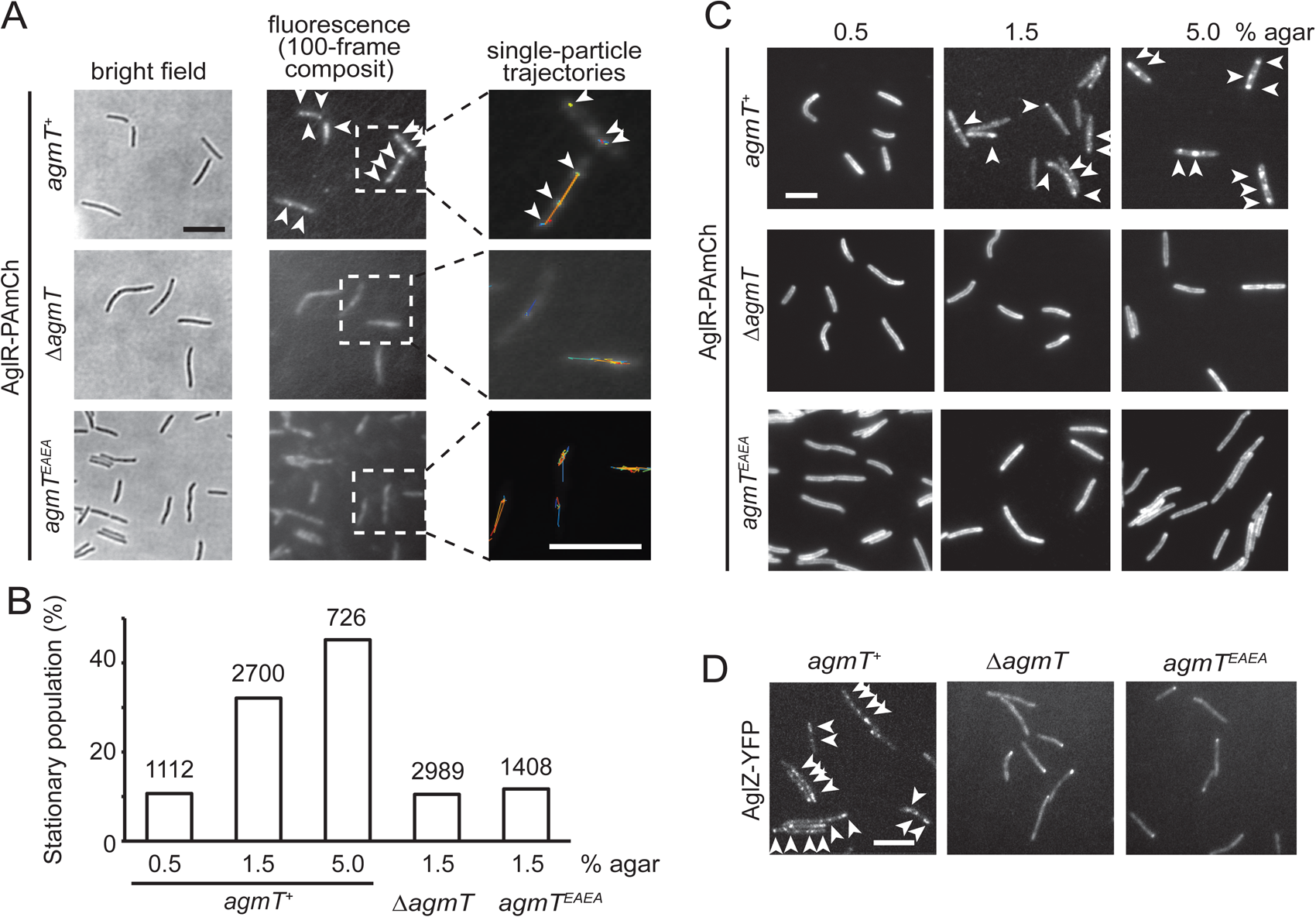
AgmT and its LTG activity are essential for proper bFAC assembly. **A)** Overall distribution of AglR-PAmCherry particles on 1.5% agar surface is displayed using the composite of 100 consecutive frames taken at 100-ms intervals. Single-particle trajectories of AglR-PAmCherry (AglR-PAmCh) were generated from the same frames. Individual trajectories are distinguished by colors. Deleting *agmT* or disabling the transglycosylase activity of AgmT (AgmT^EAEA^) decrease the stationary population of AglR particles. **B)** The stationary population of AglR-PAmCherry particles, which reflects the AglR molecules in bFACs, changes in response to substrate hardness (controlled by agar concentration) and the presence and function of AgmT. The total number of particles analyzed is shown on top of each plot. **C)** AglR fails to assemble into bFACs in the absence of active AgmT, even on 5.0% agar surfaces. **D)** AgmT and its LTG activity also support the assembly of AglZ into bFACs. White arrows point to bFACs. Scale bars, 5 µm.

bFAC assembly is sensitive to mechanical cues. As the agar concentration increases in gliding substrate, more motor molecules engage in bFACs, where they appear immotile^9,11^. We exposed *aglR-PAmCherry* cells to 405-nm excitation (0.2 kW/cm^2^) for 2 s, where most PAmCherry molecules were photoactivated and used epifluorescence to display the overall localization of AglR^11,16,30^. As the agar concentration increased, bFACs increased significantly in size (**Fig. 4C**). Accordingly, the immotile AglR particles detected by sptPALM increased from 10.7% (n = 1112) on 0.5% agar to 45.2% (n = 726) on 5.0% agar surface (**Fig. 4B**). Thus, the stationary population of AglR-PAmCherry particles reflects the AglR molecules in fully assembled bFACs.

In contrast to the cells that expressed wild-type AgmT (*agmT^+^*), AglR particles were hyper motile in the cells that lacked AgmT or expressed AgmT^EAEA^. On 1.5% agar surfaces, the immotile populations of AglR particles decreased to 10.5% (n = 2,989) and 11.7% (n = 1,408), respectively. Consistently, AglR-containing bFACs were rarely detectable in these cells, even on 5% agar surfaces (**Fig. 4A-C**).

To test if other components in the gliding machinery also depend on AgmT to assemble into bFACs, we tested the localization of AglZ, a cytoplasmic protein that is commonly used for assessing bFAC assembly^10,34^. In the cells that expressed wild-type AgmT, yellow fluorescent protein (YFP)-labeled AglZ formed a bright cluster at the leading cell pole and multiple static, near-evenly spaced clusters along the cell body, indicating the assembly of bFACs (**Fig. 4D**). In contrast, AglZ-YFP only formed single clusters at cell poles in the cells that lacked AgmT or expressed AgmT^EAEA^ (**Fig. 4D**). Taken together, the LTG activity of AgmT is essential for proper bFACs assembly.

### AgmT does not assemble into bFACs

We first hypothesized that AgmT could assemble into bFACs, where it could generate pores in PG specifically at bFAC loci that allow the inner and outer gliding complexes interact directly. Alternatively, it could bind to PG and recruit other components to bFACs through protein-protein interactions. Regardless, if AgmT assembles into bFACs, it should localize in bFACs. To test this possibility, we labeled AgmT with PAmCherry on its C-terminus and expressed the fusion protein as the sole source of AgmT using the native promoter and locus of *agmT*. *agmT-PAmCherry pilA^-^* cells displayed gliding motility that was indistinguishable from *pilA^-^* cells (**Fig. 2B**), indicating that the fusion protein is functional. Similar to many membrane proteins that resistant to dissociation by SDS^35^, immunoblot using an anti-mCherry antibody showed that AgmT-PAmCherry accumulated in two bands in SDS-PAGE that corresponded to monomers and dimers of the full-length fusion protein, respectively (**Fig. 5A**). This result is consistent with the structure of *E. coli* MltG that functions as homodimers (PDB: 2r1f). Importantly, AgmT-PAmCherry was about 50 times more abundant than AglR-PAmCherry (**Fig. 5A**). To test if AgmT assembles into bFACs, we first expressed AgmT-PAmCherry and AglZ-YFP together using their respective loci and promoter. We exposed *agmT-PAmCherry* cells to 405-nm excitation (0.2 kW/cm^2^) for 2 s to visualize the overall localization of AgmT. We found that on a 1.5% agar surface that favors gliding motility, AglZ formed bright clusters at cell poles and aggregated in near evenly-spaced bFACs along the cell body. In contrast, AgmT localized near evenly along cell bodies without forming protein clusters (**Fig. 5B**). Thus, AgmT does not localize into bFACs. These results echo the fact that despite its abundance, AgmT has not been identified as a component in bFACs despite extensive pull-down experiments using various bFAC components as baits^6,9,36^.

**Fig. 5.**
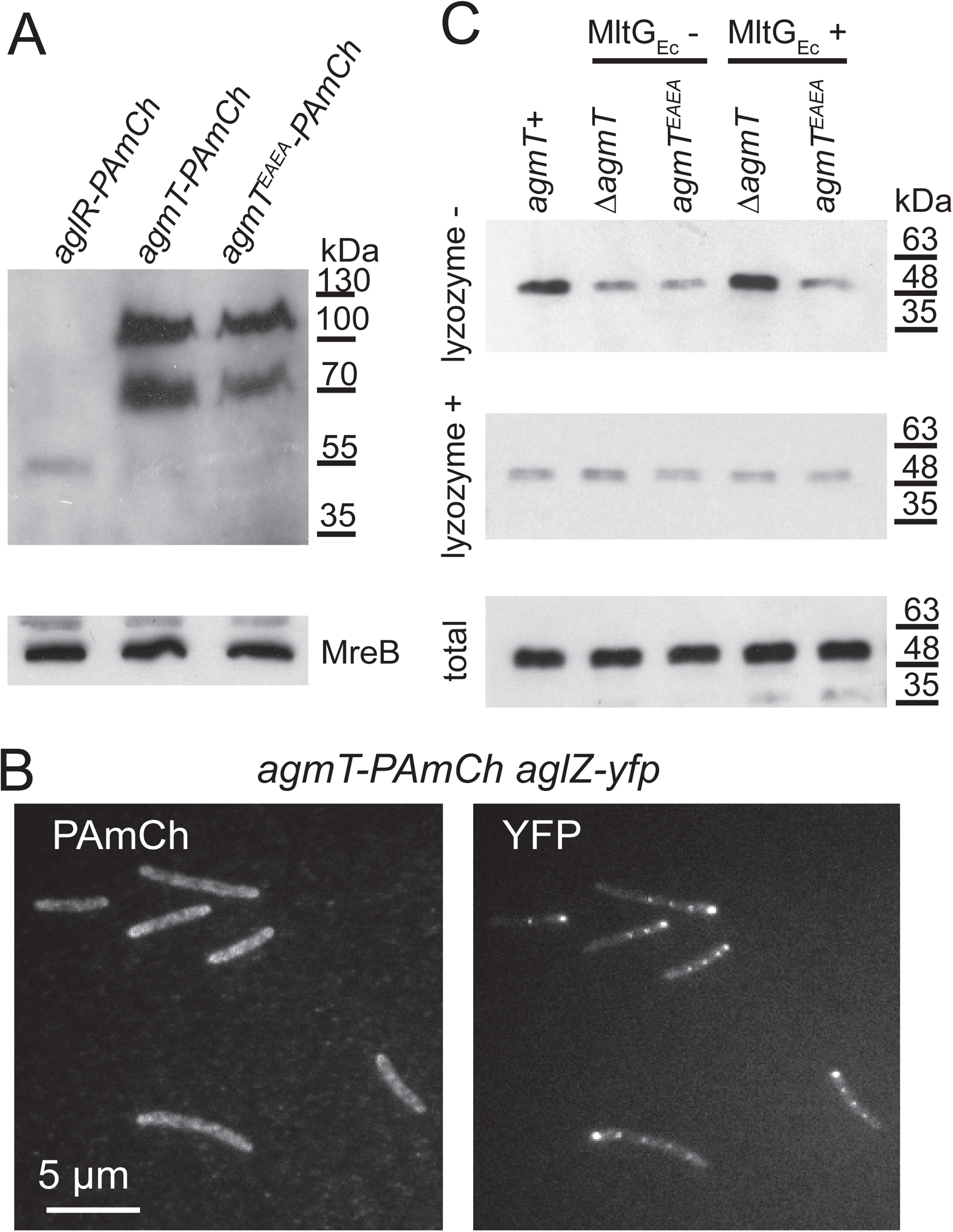
AgmT does not assemble into bFACs but connects bFACs to PG. A) Immunoblotting using *M. xanthus* cell lysates and an anti-mCherry antibody shows that PAmCherry (PAmCh)-labeled AgmT and AgmT^EAEA^ (592 amino acids) accumulate as full-length proteins. AgmT is significantly more abundant than the PAmCherry-labeled motor protein AglR (498 amino acids). The bacterial actin homolog MreB visualized using an MreB antibody is shown as a loading control. **B)** AgmT does not aggregate into bFACs. **C)** The LTG activity of AgmT is required for connecting bFACs to PG and expressing the *E. coli* LTG MltG (MltGEc) restores PG binding by bFACs in cells that lack AgmT but not in the ones that express an inactive AgmT variant (AgmT^EAEA^). AglR-PAmCherry was detected using an anti-mCherry antibody to mark the presence of bFACs that co-precipitate with PG-containing (lysozyme-) pellets. Lysates from the cells that express AglR-PAmCherry in different genetic backgrounds were pelleted by centrifugation in the presence and absence of lysozyme. The loading control of AglR-PAmCherry in the whole cell is shown as “total”.

As an additional test, we used sptPALM to track the movements of AgmT-PAmCherry single particles on 1.5 agar surfaces at 10 Hz. We reasoned that if AgmT assembles into bFACs, AgmT and gliding motors should display similar dynamic patterns. Distinct from the AglR particles of which 32.1% remained stationary, AgmT moved in a diffusive manner, showing no significant immotile population. Importantly, compared to the diffusion coefficients (*D*) of motile AglR particles (1.8 × 10^-2^ ± 3.6 × 10^-3^ µm^2^/s (n = 1,833)), AgmT particles diffused much faster (*D* = 2.9 × 10^-2^ ± 5.3 × 10^-3^ µm^2^/s (n = 8,548)). Taken together, AgmT and the gliding motor did not display significant correlation in either their localization or dynamics. We thus conclude that AgmT does not assemble into bFACs.

### AgmT connects bFACs to PG through its LTG activity

We then explored the possibility that AgmT could modify PG through its LTG activity and thus generate the anchor sites for certain components in bFACs. If this hypothesis is true, heterologous expression of a non-native LTG could rescue gliding motility in the *ΔagmT* strain. To test this hypothesis, we fused the *E. coli mltG* gene (*mltG_Ec_*) to a vanillate-inducible promoter and inserted the resulting construct to the *cuoA* locus on *M. xanthus* chromosome that does not interfere gliding^37^. We subjected the *ΔagmT* and *agmT^EAEA^* cells that expressed MltG_Ec_ to mecillinam (100 μg/ml) stress. Induced by 200 μM sodium vanillate, MltG_Ec_ restored cell morphology and integrity of the *ΔagmT* strain to the wild-type level (**Fig. 3C**). This result confirmed that MltG_Ec_ was expressed and enzymatically active in *M. xanthus*. In contrast, MltG_Ec_ failed to confer mecillinam resistance to the *agmT^EAEA^* cells (**Fig. 3C**). A potential explanation is that AgmT^EAEA^ could still bind to PG and thus block MltG_Ec_ from accessing *M. xanthus* PG.

Consistent with its LTG activity, the expression of MltG_Ec_ restored gliding motility of the *ΔagmT pilA^-^* cells on both the colony (**Fig. 2B**) and single-cell (**Fig. 2C, D**) levels. Interestingly, in the absence of sodium vanillate, the leakage expression of MltG_Ec_ using the vanillate-inducible promoter was sufficient to compensate the loss of AgmT. A plausible explanation of this observation is that as *E. coli* grows much faster (generation time 20 - 30 min) than *M. xanthus* (generation time ∼4 h), MltG_Ec_ could possess significantly higher LTG activity than AgmT. Induced by 200 µM sodium vanillate, the expression of MltG_Ec_ further but non significantly increased the velocity and gliding persistency (**Fig. 2B-D**). Importantly, the expression of MltG_Ec_ failed to restore gliding motility in the *agmT^EAEA^ pilA* cells, even in the presence of 200 µM sodium vanillate (**Fig. 2B**). Consistent with the mecillinam resistance assay (**Fig. 3C**), this result suggests that AgmT^EAEA^ still binds to PG and that in the absence of its LTG activity, AgmT does not anchor bFACs to PG. As MltG_Ec_ substitutes AgmT in gliding motility, it is the LTG activity, rather than specific interactions between AgmT and bFACs, that is required for gliding.

It is challenging to visualize how AgmT facilitates bFAC assembly through its enzymatic activity *in vitro*. First, the inner complex in a bFAC spans the cytoplasm, inner membrane, and periplasm and contains multiple proteins whose activities are interdependent^8–10^. It is hence difficult to purify the entire inner complex in its functional state. Second, Faure *et al*. hypothesized that AglQ and AglS in the gliding motor could bind PG directly^10^. However, such binding has not been proved experimentally due to the difficulty in purifying the membrane integral AglRQS complex. To overcome these difficulties, we used AglR-PAmCherry to represent the inner complex of bFAC and investigated how bFACs bind PG in their native environment. To do so, we expressed AglR-PAmCherry in different genetic backgrounds as the sole source of AglR using the native *aglR* locus and promoter. We lysed these cells using sonication, subjected their lysates to centrifugation, isolated the pellets that contained PG, and detected the presence of AglR-PAmCherry in these pellets by immunoblots using an mCherry antibody. Eliminating AgmT and disabling its active site significantly reduced the amounts of AglR in the pellets (**Fig. 5C**). Because such pellets contain both the PG and membrane fractions, we further eliminated PG in cell lysates using lysozyme before centrifugation and determined the amount of AglR-PAmCherry in the pellets that only contained membrane fractions. Regardless of the presence and activity of AgmT, comparable amounts of AglR precipitated in centrifugation pellets after lysozyme treatment, indicating that AgmT does not affect the expression or stability of AglR (**Fig. 5C**). Thus, active AgmT facilitates the association between bFACs and PG. Strikingly, the heterologous expression of MltG_Ec_ enriched AglR-PAmCherry in the PG-containing pellets from the *ΔagmT* cells but not the *agmT^EAEA^* ones (**Fig. 5C**). These results indicate that AgmT connects bFACs to PG through its LTG activity.

The assembly of bFACs produces wave-like deformation on cell surface^6,38^, suggesting that their assembly may require a flexible PG layer^2,6,11,12^. As a major contributor to cell stiffness, PG flexibility affects the overall stiffness of cells^39^. To test the possibility that AgmT and MltG_Ec_ facilitate bFAC assembly by reducing PG stiffness, we adopted the GRABS assay^39^ to quantify if the lack of AgmT and the expression of MltG_Ec_ affects cell stiffness. To quantify changes in cell stiffness, we simultaneously measured the growth of the *pilA^-^*, *ΔagmT pilA^-^*, and *ΔagmT P_van_-MltG_Ec_ pilA^-^* (with 200 µM sodium vanillate) cells in a 1% agarose gel infused with CYE and liquid CYE and calculated the GRABS scores of the *ΔagmT pilA^-^*, and *ΔagmT P_van_-MltG_Ec_ pilA^-^* cells using the *pilA^-^* cells as the reference, where positive and negative GRABS scores indicate increased and decreased stiffness, respectively (see Materials and Methods and Ref^39^). The GRABS scores of the *ΔagmT pilA^-^*, and *ΔagmT P_van_-MltG_Ec_ pilA^-^* (with 200 µM sodium vanillate) cells were -0.06 ± 0.04 and -0.10 ± 0.07 (n = 4), respectively, indicating that neither AgmT nor MltG_Ec_ affects cell stiffness significantly. Whereas PG flexibility could still be essential for gliding, AgmT and MltG_Ec_ do not regulate bFAC assembly by modulating PG stiffness. Instead, these LTGs could connect bFACs to PG by generating structural features that are irrelevant to PG stiffness.

## Discussion

Gliding bacteria adopt a broad spectrum of nanomachineries. Compared to the trans-envelope secretion system that drives gliding in *Flavobacterium johnsoniae* and *Capnocytophaga gingivalis*, the gliding machinery of *M. xanthus* appears to lack a stable structure that transverses the cell envelope, especially across the PG layer^2,40,41^. In order to generate mechanical force from the fluid *M. xanthus* gliding machinery, the inner complex must establish solid contact with PG and push the outer complex to slide^10,19^. In this work, we discovered AgmT as the long-sought factor that facilitates persistent gliding by connecting bFACs to PG.

It is surprising that AgmT itself does not assemble into bFACs and that MltG_Ec_ substitutes AgmT in gliding. Thus, rather than interacting with bFAC components directly and specifically, AgmT facilitates proper bFAC assembly indirectly through its LTG activity. LTGs usually break glycan strands and produce unique anhydro caps on their ends^42–46^. However, because AgmT is the only LTGs that is required for gliding, it is not likely to facilitate bFAC assembly by generating such modification on glycan strands. *E. coli* MltG is a glycan terminase that controls the length of newly synthesized PG glycans^26^. Likewise, AgmT could generate short glycan strands and thus uniquely modify the overall structure of *M. xanthus* PG, such as producing small pores that retard and retain the inner subcomplexes of bFACs (**Fig. 6**). On the contrary, the *M. xanthus* mutants that lack active AgmT could produce PG with increased strain length, which blocks bFACs from binding to the cell wall and precludes stable bFAC assembly. However, it would be very difficult to demonstrate how glycan length affects the connection between bFACs and PG.

**Fig. 6.**
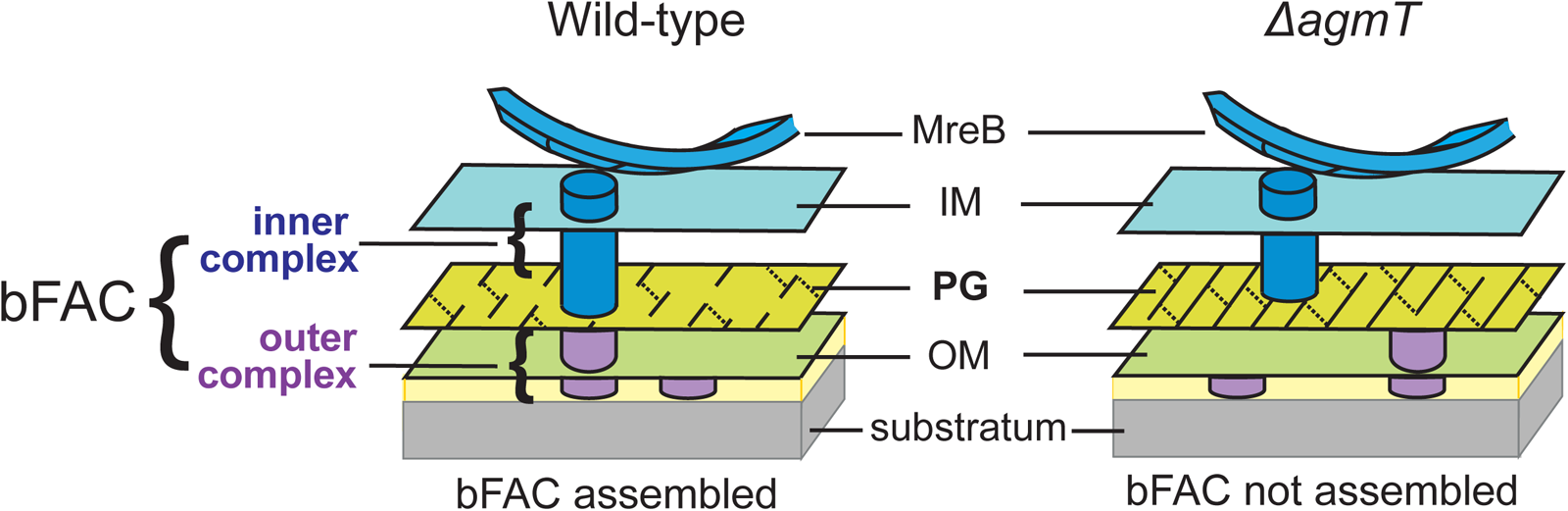
A possible mechanism by which AgmT connect bFACs to PG. AgmT could generate short glycan strands through its LTG activity and thus uniquely modify the overall structure of *M. xanthus* PG, such as producing small pores that retard and retain the inner subcomplexes of bFACs. Likewise, the *M. xanthus* mutants that lack active AgmT could produce PG with increased strain length, which precludes bFACs from binding to the cell wall.

Then how does AgmT, a protein localized diffusively, facilitate bFAC assembly into near-evenly spaced foci? The actin homolog MreB is the only protein in bFACs that displays quasiperiodic localization independent of other components^16,21^. *M. xanthus* MreB assembles into bFACs and positions the latter near evenly along the cell body^6,11,16,21–23^. MreB filaments change their orientation in accordance with local membrane curvatures and could hence respond to mechanical cues^47–49^. Strikingly, *M. xanthus* does assemble bFACs in response to substrate hardness (**Fig. 4C**) and assembled bFACs could distort the cell envelope, generating undulations on the cell surface^6,9,11,38,50^. Taken together, whereas AgmT potentially modifies the entire PG layer, it is still the quasiperiodic MreB filaments that determine the loci for bFAC assembly in response to the mechanical cues from the gliding substrate (**Fig. 6**).

Most bacteria encode multiple LTGs that function in PG growth, remodeling, and recycling^43,46^. This work provides an example that macro bacterial machineries domesticate non-specialized LTGs for specialized functions. Other examples include MltD in the *Helicobacter pylori* flagellum and MltE in the *E. coli* type VI secretion system^51,52^. Based on their catalytic folds and domain arrangements, LTGs can be categorized inti six distinct families^43^. Both *M. xanthus* AgmT and *E. coli* MltG belong to the YceG/MltG family, which is the first identified LTG family that is conserved in both Gram-negative and positive bacteria^26,43^. About 70% of bacterial genomes, including firmicutes, proteobacteria, and actinobacteria, encode YceG/MltG domains^26^. The unique inner membrane localization of this family and the fact that AgmT is the only *M. xanthus* LTG that belongs to this family (**Table S2**) could partially explain why it is the only LTG that contributes to gliding motility. It will be interesting to investigate if other LTGs, once anchored to the inner membrane, could also facilitate force-generation by bFACs.

## Methods

### Strains and growth conditions

*M. xanthus* strains used in this study are listed in **Table S2**. Vegetative *M. xanthus* cells were grown in liquid CYE medium (10 mM MOPS pH 7.6, 1% (w/v) Bacto™ casitone (BD Biosciences), 0.5% yeast extract and 8 mM MgSO_4_) at 32 °C, in 125-ml flasks with vigorous shaking, or on CYE plates that contains 1.5% agar. All genetic modifications on *M. xanthus* were made on the chromosome. Deletion and insertion mutants were constructed by electroporating *M. xanthus* cells with 4 µg of plasmid DNA. Transformed cells were plated on CYE plates supplemented with 100 µg/ml sodium kanamycin sulfate and 10 µg/ml tetracycline hydrochloride when needed. AgmT and AgmT^EAEA^ were labeled with PAmCherry at their C-termini by fusing their gene to a DNA sequence that encodes PAmCherry through a KESGSVSSEQLAQFRSLD (AAGGAGTCCGGCTCCGTGTCCTCCGAGCAGCTGGCCCAGTTCCGCTCCCTGGAC) linker. All constructs were confirmed by PCR and DNA sequencing.

### Immunoblotting

The expression and stability of PAmCherry-labeled proteins were determined by immunoblotting following SDS-PAGE using an anti-mCherry antibody (Rockland Immunochemicals, Inc., Lot 46705) and a goat anti-Rabbit IgG (H+L) secondary antibody, HRP (Thermo Fisher Scientific, catalog # 31460). MreB was detected as the loading control using an anti-MreB serum^21^ and the same secondary antibody. The blots were developed with Pierce™ ECL Western Blotting Substrate (Thermo Fisher Scientific REF 32109) and a MINI-MED 90 processor (AFP Manufacturing).

### Gliding assay

Five microliters of cells from overnight culture were spotted on CYE plates containing 1.5% agar at 4 ×10^9^ colony formation units (cfu)/ml and incubated at 32 °C for 48 h. Colony edges were photographed using a Nikon Eclipse™ e600 phase-contrast microscope with a 10× 0.30 NA objective and an OMAX™ A3590U camera.

### Protein expression and purification

DNA sequences encoding amino acids 25 – 339 of AgmT and AgmT^EAEA^ were amplified by polymerase chain reaction (PCR) and inserted into the pET28a vector (Novogen) between the restriction sites of *Eco*RI and *Hind*III and used to transform *E. coli* strain BL21(DE3). Transformed cells were cultured in 20 ml LB (Luria-Bertani) broth at 37 °C overnight and used to inoculate 1 L LB medium supplemented with 1.0% glucose. Protein expression was induced by 0.1 mM IPTG (isopropyl-h-d-thiogalactopyranoside) when the culture reached an OD_600_ of 0.8. Cultivation was continued at 16 °C for 10 h before the cells were harvested by centrifugation at 6,000 × g for 20 min. Proteins were purified using a NGC™ Chromatography System (BIO-RAD) and 5-ml HisTrap™ columns (Cytiva)^15,53^. Purified proteins were concentrated using Amicon™ Ultra centrifugal filter units (Millipore Sigma) with a 10-kDa molecular weight cutoff and stored at -80 °C.

### LTG activity (RBB) assay

PG was purified following the published protocol^30,54^. In brief, *M. xanthus* cells were grown until mid-stationary phase and harvested by centrifugation (6,000 × g, 20 min, 25 °C). Supernatant was discarded and the pellet was resuspended and boiled in 1× PBS with 5% SDS for 2 h. SDS was removed by repetitive wash with water and centrifugation (21,000 × g, 10 min, 25 °C). Purified PG from 100 ml culture was suspended into 1 ml 1× PBS and stored at -20 °C. RBB labelling of PG was performed essentially as previously described^28,29^. Purified sacculi were incubated with 20 mM RBB in 0.25 M NaOH overnight at 37 °C. Reactions were neutralized by adding equal volumes of 0.25 M HCl and RBB-labeled PG was collected by centrifugation at 21,000 × g for 15 min. Pellets were washed repeatedly with water until the supernatants became colorless. RBB-labelled sacculi were incubated with purified AgmT and AgmT^EAEA^ (1 mg/ml) at 25 °C for 12 h. lysozyme (1 mg/ml) was used as a positive control. Dye release was quantified by the absorption at 595 nm from the supernatants after centrifugation (21,000 × g, 10 min, 25 °C).

### Cell stiffness (GRABS) assay

Cell stiffness was quantified using the GRABS assay^55^. Briefly, overnight cultures of *pilA^-^*, *ΔagmT pilA^-^*, and *ΔagmT P_van_-MltG_Ec_ pilA^-^* (with 200 µM sodium vanillate) cells were inoculated to liquid CYE medium or embedded in solid CYE medium with 1% agarose to OD_600_ 0.5 in 2-ml optical spectrometer cuvettes and incubated at 32 °C. Growth data were collected over 24 hr, and the GRABS scores were calculated as (OD_mutant, agarose_/OD*_pilA_*_, agarose_) - (OD_mutant, liquid_/OD*_pilA_*_, liquid_).

### Co-precipitation assay

*M. xanthus* cells expressing AglR-PAmCherry were grown in liquid CYE to OD_600_ ∼1, harvested by centrifugation (6,000 × g, 20 min, 25 °C), washed by 1× PBS, and resuspended into 1× PBS to OD_600_ 6. Cells (1 ml) were lysed using a Cole-Parmer 4710 Ultrasonic Homogenizer. Unbroken cells and large debris were eliminated by centrifugation (6,000 × g, 10 min, 25 °C). Supernatants were subjected to centrifugation at 21,000 × g for 15 min. Pellets that contain both the PG and membrane fractions were resuspended to 1 ml in 1× PBS. To collect the pellets that do not contain PG, supernatants from sonication lysates were incubated with 5 mg/ml lysozyme at 25 °C for 5 h before centrifugation. Five microliters of each resuspended pellet were mixed with 5 µl 2× loading buffer and applied to SDS-PAGE. AglR-PAmCherry was detected using immunoblotting.

### Microscopy Analysis

For all imaging experiments, we spotted 5 μl of cells grown in liquid CYE medium to OD_600_ ∼1 on agar (1.5%) pads. For the treatments with inhibitors, inhibitors were added into both the cell suspension and agar pads. The length and width of cells were determined from differential interference contrast (DIC) images using a MATLAB (MathWorks) script^16,30,56^. DIC and fluorescence images of cells were captured using an Andor iXon Ultra 897 EMCCD camera (effective pixel size 160 nm) on an inverted Nikon Eclipse-Ti™ microscope with a 100× 1.49 NA TIRF objective. For sptPALM, *M. xanthus* cells were grown in CYE to 4 ×10^8^ cfu/ml and PAmCherry was activated using a 405-nm laser (0.3 kW/cm^2^), excited and imaged using a 561-nm laser (0.2 kW/cm^2^). Images were acquired at 10 Hz. For each sptPALM experiment, single PAmCherry particles were localized in at least 100 individual cells from three biological replicates. sptPALM data were analyzed using a MATLAB (MathWorks) script^16,56^. Briefly, cells were identified using differential interference contrast images. Single PAmCherry particles inside cells were fit by a symmetric 2D Gaussian function, whose center was assumed to be the particle’s position^16^. Particles that explored areas smaller than 160 nm × 160 nm (within one pixel) in 0.4 - 1.2 s were considered immotile^16,30,56^. Sample trajectories were generated using the TrackMate^57^ plugin in the ImageJ suite (https://imagej.net).

## Acknowledgements

We thank the Department of Biology and the College of Arts and Sciences at Texas A&M University for the support on camera purchase. This work is supported by the National Institute of Health R01GM129000 to B. N.. C. A. R. C. was supported in the 2023 – 2024 academic year by a National Institute of Health Diverse Predoctoral Training in Genetics grant T32GM135115 awarded to the Genetics and Genomics Interdisciplinary Program at Texas A&M University.

## Author Contributions

C. R. C., O. G. F., and B. N. designed the study, performed experiments and data analysis, and wrote the manuscript. All authors read and approved the manuscript.

## Data availability

All data generated or analyzed during this study are included in the manuscript and supporting files.

