## Supplementary info for "A lytic transglycosylase connects bacterial focal adhesion complexes to the peptidoglycan cell wall"

[§]C. A. R. C. and O. G. F. contribute equally to this work

*Corresponding author: Beiyan Nan, 306C BSBE, 3258 TAMU, College Station, TX77843

### Table S1. primers used to knock out the genes that encode potential LTGs

| ORF tag in DZ2 genome[1] (LTG family) | Corresponding ORF in DK1622 genome[2] | Primers for plasmid-insertion knockout | Primers for in-frame deletion |
|---|---|---|---|
| K1515_37860 (MltE) | MXAN_RS00570 | CCCAAGCTTCCCCATGCGCCTCTCTCCTC<br>CGGGATCCAGCGCCGGAGGCAGGTTGCC | N/A |
| K1515_37385 (MltE) | MXAN_RS01035 | CCCAAGCTTTCCGAGGACCAGTTGGAAGC<br>CGGGATCCTCGTTCGACAGCGGACTGGTG | N/A |
| K1515_34725 (unknown) | MXAN_RS03640 | CCCAAGCTTGTTCATGCTGGGCCTGGCCG<br>CGGGATCCTTCTGCATGAAGGTGTGGAT | N/A |
| K1515_24775 (RlpA) | MXAN_RS12365 | N/A (too short for plasmid-insertion) | CCCAAGCTTCGTCAAGTTCTCCTCGAAGCT<br>CGGGATCCTAGCGGGTGATGTCGTTGGTCA<br>GGGGTACCGCGCCGGTGTAGTGAGCCGTC<br>CGAATTCCGGAAAGCGCGCCAGTTGCG |
| K1515_22185 (MltE) | MXAN_RS14935 | CCCAAGCTTCGGGCGGGAGGTCTTTCGG<br>CGGGATCCTGTAACGAGCGCAGATAACG | N/A |
| K1515_20905 (MltE) | MXAN_RS16205 | CCCAAGCTTCAGAGGCGCAGTTGTCGCCC<br>CGGGATCCACCAGCGCCTCGCCACGCCCC | N/A |
| K1515_20820 (MltE) | MXAN_RS16290 | CCCAAGCTTCAGTACATCCGTACGAGATGA<br>CGGGATCCGCCAGCCAGGTGCCGGTGTC | N/A |
| K1515_17460 (MltE) | MXAN_RS19615 | CCCAAGCTTGCGGGGATGTCGCTGGGACA<br>CGGGATCCTTGTCGAGCTTCACCTTGTC | N/A |
| K1515_14545 (MltE) | MXAN_RS22465 | CCCAAGCTTGTCGGCGGGGGCGGATGCC<br>CGGGATCCGGCGTTGTACGCGGCAACCAT | N/A |
| K1515_09355 (MltD/E) | MXAN_RS27580 | CCCAAGCTTCCTGCCAAGGCTCAGGTGGT<br>CGGGATCCTGCTTCAGCCCGTACTGCTTG | N/A |
| K1515_06035 (MltE) | MXAN_RS30855 | CCCAAGCTTGGGGGGATGACGGTGCCCGG<br>CGGGATCCATTCACGGAGCCTGGGCCCGC | N/A |
| **K1515_04910 (AgmT/MltG/YceG)** | MXAN_RS31980 | N/A | CCCAAGCTTTGCTGGCGCCCTTCGTCG<br>CGGGATCCTCGCGGCGCACGCTATCAC<br>CGGGGTACCGGCATGAACTCAGCGCCC<br>CGGAATTCACGATACCGCCCGACGCAA |
| K1515_01490 (MltA) | MXAN_RS35365 | CCCAAGCTTCTCGCCTCGGCCTGTCCGA<br>CGGGATCCTTGGGGATGGCGCCCTCCTG | N/A |
| K1515_01440 (MltD/E) | MXAN_RS35415 | CCCAAGCTTGTCGGCGCGGCAGTCCTTGA<br>CGGGATCCGGGTGGTGGGATGGGACATG | N/A |

**Table S2. *M. xanthus* strains used in this work**

| *M. xanthus* strains | Source | Identifier |
|---|---|---|
| DZ2 (wild-type *M. xanthus* strain) | [3] | DZ2 |
| *pilA::tet* | [4] | DZ4469 |
| *aglR-PAmCherry* | [5] | N/A |
| *Δ3 (Δpbp1a1 Δpbp1a2 pbp1c::kan)* | [6] | BN311 |
| *aglZ-yfp::kan* | [7] | TM7 |
| *Δ3 (Δpbp1a1 Δpbp1a2 pbp1c::kan) pilA::tet* | This study | BN328 |
| *ΔagmT* | This study | BN329 |
| *ΔagmT pilA::tet* | This study | BN330 |
| $agmT^{EAEA}$ | This study | BN331 |
| $agmT^{EAEA}$ *pilA::tet* | This study | BN332 |
| *ΔaglR pilA::tet* | This study | BN333 |
| *agmT-PAmCherry* | This study | BN334 |
| $agmT^{EAEA}$*-PAmCherry* | This study | BN335 |
| *agmT-PAmCherry pilA::tet* | This study | BN336 |
| *ΔagmT pilA::tet pMR3679* ($P_{van}$ vector control) | This study | BN337 |
| *ΔagmT pMR3679-*$mltG_{EC}$ | This study | BN338 |
| *ΔagmT pMR3679-*$mltG_{EC}$ *pilA::tet* | This study | BN339 |
| $agmT^{EAEA}$ *pMR3679-*$mltG_{EC}$ | This study | BN340 |
| $agmT^{EAEA}$ *pMR3679-*$mltG_{EC}$ *pilA::tet* | This study | BN341 |
| *aglR-PAmCherry ΔagmT* | This study | BN342 |
| *aglR-PAmCherry* $agmT^{EAEA}$ | This study | BN343 |
| *aglZ-yfp::kan ΔagmT* | This study | BN344 |
| *aglZ-yfp::kan* $agmT^{EAEA}$ | This study | BN345 |
| *aglR-PAmCherry aglZ-yfp::kan* | This study | BN346 |
| *K1515_37860::kan pilA::tet* | This study | BN347 |
| *K1515_37385::kan pilA::tet* | This study | BN348 |
| *K1515_34725::kan pilA::tet* | This study | BN349 |
| *ΔK1515_24775 pilA::tet* | This study | BN350 |
| *K1515_22185::kan pilA::tet* | This study | BN351 |
| *K1515_20905::kan pilA::tet* | This study | BN352 |
| *K1515_20820::kan pilA::tet* | This study | BN353 |
| *K1515_17460::kan pilA::tet* | This study | BN354 |
| *K1515_14545::kan pilA::tet* | This study | BN355 |
| *K1515_09355::kan pilA::tet* | This study | BN356 |
| *K1515_06035::kan pilA::tet* | This study | BN357 |
| *K1515_01490::kan pilA::tet* | This study | BN358 |
| *K1515_01440::kan pilA::tet* | This study | BN359 |

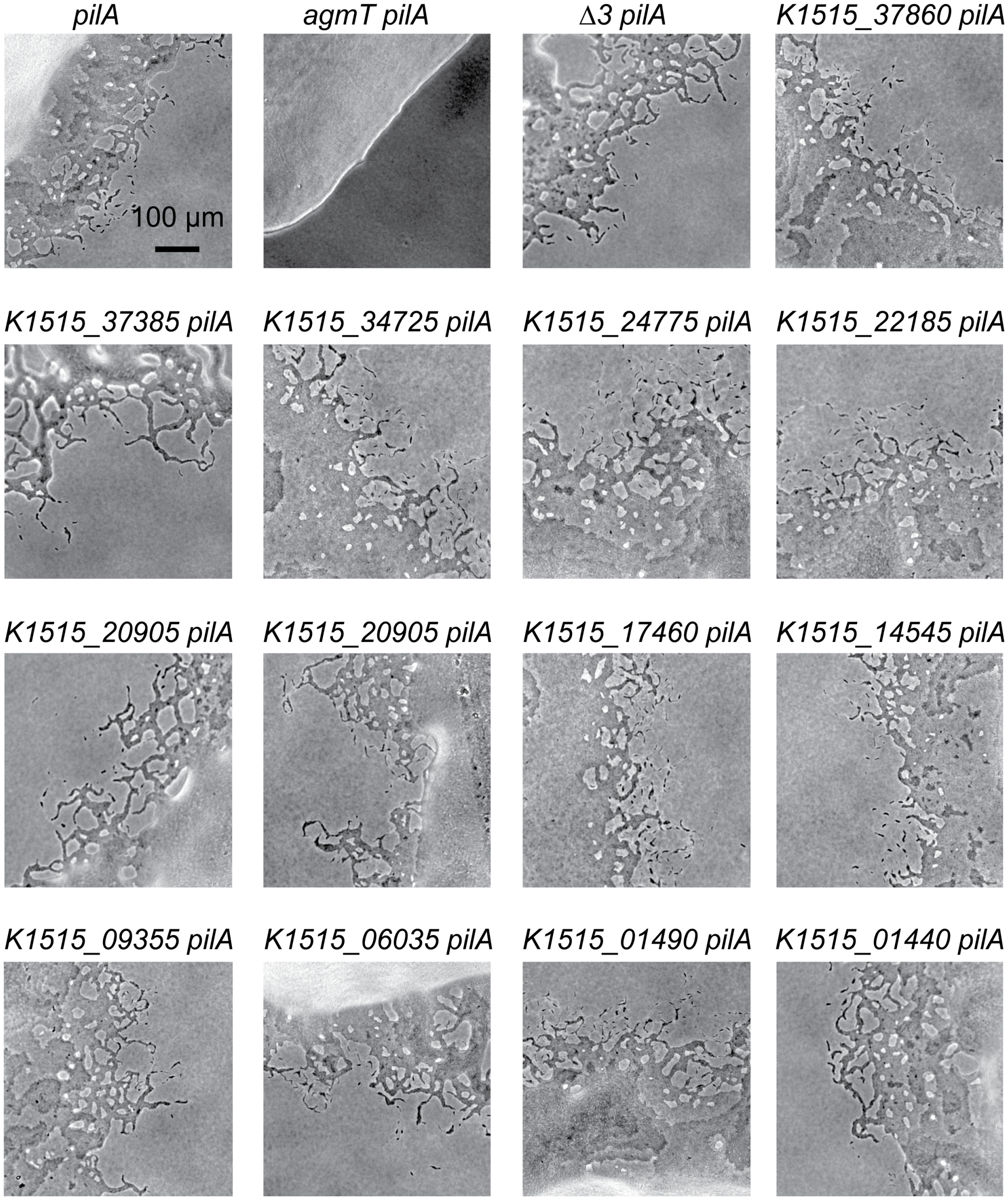


**Fig. S1. AgmT is the only LTG that is required for *M. xanthus* gliding motility.** All 14 genes that encode putative LTGs were knocked out individually in the *pilA* background. Single cells moving out from colony edges display functional gliding motility. Colony edges were imaged after incubating cells on 1.5% agar surfaces for 24 h.

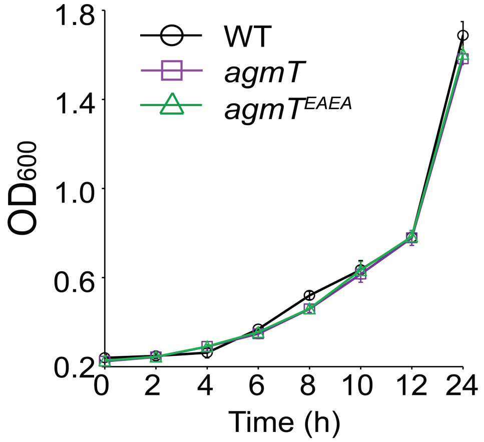


**Fig. S2. AgmT does not regulate growth.** Cells were inoculated at $OD_{600}$ 0.1 and growth measured for 24 h from three biological replicates. Data are presented as mean values ± SD.

**Videos**

**Video 1. Gilding motility of the *ΔagmT pilA*⁻ cells.** Timelapse was captured at 10-s intervals and the video plays at 10 frames/s (100 X speedup).

**Video 2. Gilding motility of the *pilA*⁻ cells.** Timelapse was captured at 10-s intervals and the video plays at 10 frames/s (100 X speedup).
